# SeQuiLa-cov: A fast and scalable library for depth of coverage calculations

**DOI:** 10.1101/494468

**Authors:** Marek Wiewiórka, Agnieszka Szmurło, Wiktor Kuśmirek, Tomasz Gambin

**Author notes:** Contributed equally (Marek Wiewiórka, Agnieszka Szmurło).

## Abstract

**Background:** Depth of coverage calculation is an important and computationally intensive preprocessing step in a variety of next generation sequencing pipelines, including the analyses of RNA-seq data, detection of copy number variants, or quality control procedures.

**Results:** Building upon big data technologies, we have developed SeQuiLa-cov, an extension to the recently released SeQuiLa platform, which provides efficient depth of coverage calculations, reaching more than 100x speedup over the state-of-the-art tools. Performance and scalability of our solution allows for exome and genome-wide calculations running locally or on a cluster while hiding the complexity of the distributed computing with Structured Query Language Application Programming Interface.

**Conclusions:** SeQuiLa-cov provides significant performance gain in depth of coverage calculations streamlining the widely used bioinformatic processing pipelines.

## Introduction

Given a set of sequencing reads and a genomic contig, depth of coverage for a given position is defined as a total number of reads overlapping the locus.

The coverage calculation is a frequently performed but time-consuming step in the analysis of Next Generation Sequencing (NGS) data. In particular, Copy-Number Variant detection pipelines require obtaining sufficient read depth of the analyzed samples [1, 2, 3]. In other applications, the coverage is computed to assess the quality of the sequencing data (e.g. to calculate the percentage of genome with at least 30X read depth) or to identify genomic regions overlapped by insufficient number of reads for reliable variant calling [4]. Finally, depth of coverage is one of the most computationally intensive parts of differential expression analysis using RNA-seq data at single-base resolution [5, 6, 7].

A number of tools supporting this operation have been developed, with 22 of them specified in Omictools catalog [8]. Well known, state-of-the-art solutions include: samtools *depth* [9], bedtools *genomecov* [10], GATK *DepthOfCoverage* [11], sambamba [12], and mosdepth [13] (see comparison presented in Table 1).

**Table 1.** Comparison of leading coverage calculation software tools.

| tool | approach | Functionality |  |  | language | Implementation |  |  |
| --- | --- | --- | --- | --- | --- | --- | --- | --- |
|  |  | bases | blocks | windows |  | Intel GKL | parallelism type | interface |
| samtools | pileup | yes | no | no | C | no | none | cmd line |
| bedtools | events | yes | yes | no | C++ | no | none | cmd line |
| GATK<sup>1</sup> | pileup | yes | no | no | Java | yes | distributed | cmd line |
| sambamba | pileup | no | yes | yes | D | no | multithreaded | cmd line |
| mosdepth | events | no | yes | yes | Nim | no | multithreaded<sup>2</sup> | cmd line |
| SeQuiLa-cov | events | yes | yes | yes | Scala | yes | distributed | Scala, SQL |

<sup>1</sup>GATK *DepthOfCoverage* has not yet been ported to the latest version, i.e. GATK 4.x
<sup>2</sup>Only for BAM decompression

Traditionally, these methods calculate the depth of coverage using a pileup-based approach (introduced in samtools [9] and used in GATK [11]), which is inefficient since it iterates through each nucleotide position at every read in a Binary Alignment Map (BAM) file. An optimized, event-based approach has been proposed in bedtools [10] and mosdepth [13]. These algorithms use only specific ‘events’, i.e. start and end of the alignment blocks within each read (Figure 1A) instead of analyzing every base of each read, which significantly reduces the overall computational complexity.

**Figure 1.**
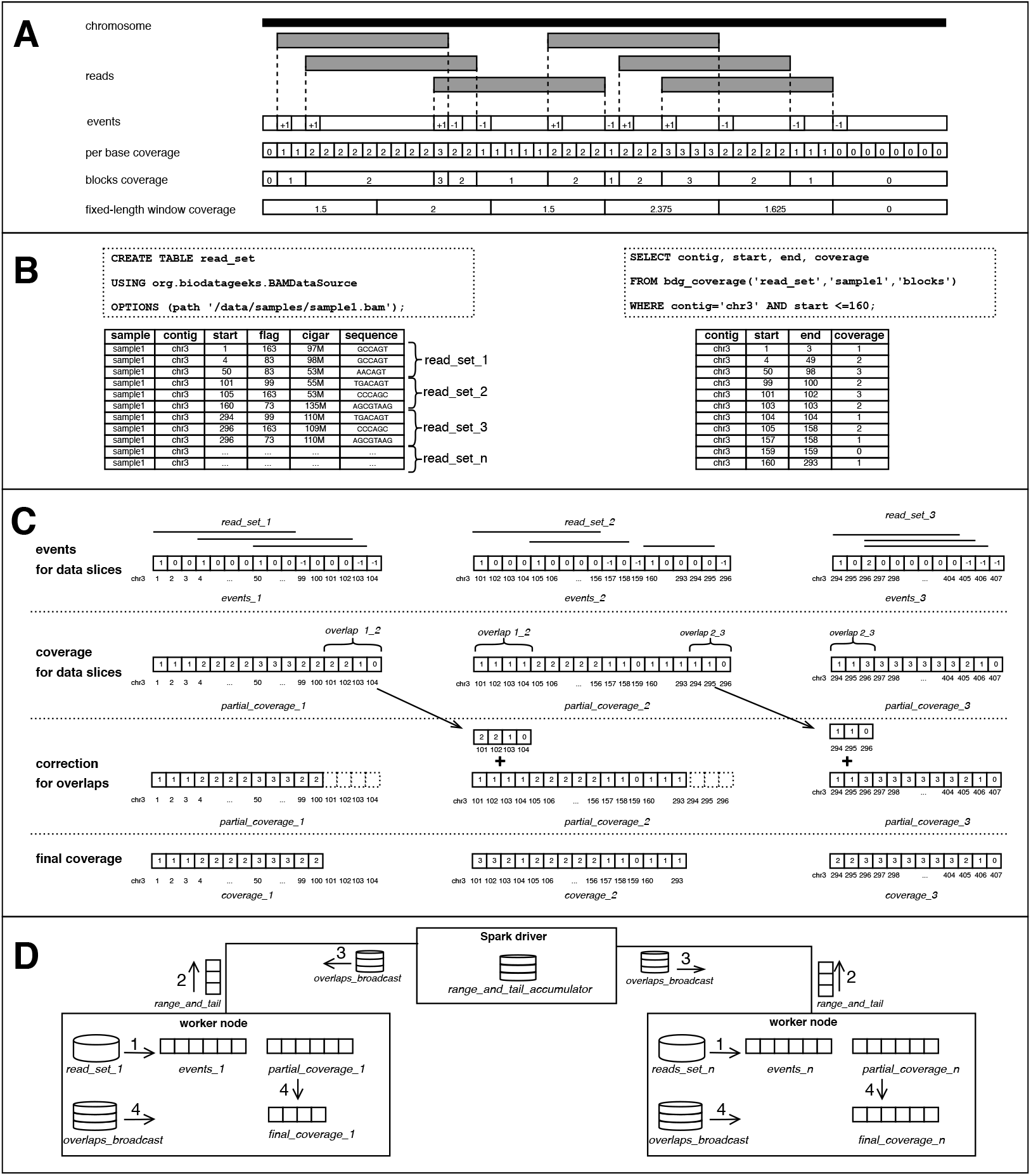
SeQuiLa-cov: functionality, algorithm and implementation. **Panel A** shows the general concept of events-based algorithm for depth of coverage calculation. Given a genomic chromosome and a set of aligned sequencing reads, the algorithm allocates *events* vector. Subsequently, it iterates the list of reads and increments/decrements by one the values of the *events* vector at the indexes corresponding to start/end positions of each read. The depth of coverage for a genomic locus is calculated using the cumulative sum of all elements in the *events* vector preceding specified position. The algorithm may produce three typically used coverage types: (i) *per-base* coverage, which includes the coverage value for each genomic position separately, (ii) *blocks* which lists adjacent positions with equal coverage values are merged into single interval, and (iii)*fixed-length windows* coverage that generates set of equal-size, non-overlapping and tiling genomic ranges and outputs arithmetic mean of base coverage values for each region. **Panel B** presents the provided SQL API to interact with NGS data. The first statement creates a relational table *read_set* over compressed BAM files using the provided custom Data Source, whereas the second statement demonstrates the use of *bdg_coverage* function to calculate depth of coverage for a specified sample. The presented call for coverage method takes sample identifier (sample1) and result type (blocks) as input parameters. *bdg_coverage* is implemented as a tablevalued function. Therefore, it outputs a table as a result allowing for customizing a query using Data Manipulation Language e.g. in the SELECT or WHERE clause. For the purpose of this example, we assume that BAM file for sample1 contains only reads from chr3. **Panel C** shows the concept of distributed version of events-based algorithm. Assuming that we run our calculations in a distributed environment, the computation nodes do not work on the whole input data set (table *read_set)* but on *n* smaller data partitions (*slice*_1_, *slice*_2_, …, *slice_n_*), each containing subset of input aligned reads. First the algorithm calculates partial *events* vector for available data slices and subsequently produces corresponding partial *partial_coverage* vector. Due to the possibility of overlapping of ranges between two consecutive data slices, additional correction step needs to be performed. When an overlap is identified, the corresponding coverage values from the preceding vector’s tail are cut and added to the head values of the subsequent vector. On the figure two overlaps were shown, one of them situated between *partial_coverage_1_* and *partial_coverage_2_* (*overlap*_12_ of length 4) encompassing positions chr3:101-104. The coverage values from *partial_coverage*_1_ for *overlap*_12_ are removed from *partial_coverage*_1_ and added to the head of *partial_coverage*_2_. As a result, a set of non-overlapping coverage vectors are calculated, which is further integrated into the depth of coverage for the whole input data set. **Panel D** presents the implementation details of SeQuiLa-cov. We have used the Apache Spark environment, where a single driver node runs the high-level driver program, which schedules tasks for multiple worker nodes. On each worker node, a set of data partitions are accessed and manipulated in order to generate *events* and *partial_coverage* vectors. To gather data about *partial_coverage* vectors’ ranges along with tailing coverage values, and to distribute data needed for rearranging coverage vector values and ranges, we have used Spark’s shared variables *accumulator* and *broadcast*, respectively.

Samtools and bedtools depth of coverage modules do not provide any support for multi-core environment. Mosdepth implements parallel BAM decompression, but its main algorithm remains sequential. Sambamba, on the other hand, promotes itself as a highly parallel tool, implementing depth of coverage calculations in a map-reduce fashion utilizing multiple threads on a single node. Regardless of parallelization degree, all of the above mentioned tools share a common bottleneck caused by using a single thread for returning results. Finally, GATK was the first genomic framework providing a support for distributed computations, however, the *DepthOfCoverage* method has not been ported yet to the current software release of the toolkit.

#### Key Points

- SeQuiLa-cov allows for high-coverage (~60x) genome-wide depth of coverage calculations in less than one minute.
- SeQuiLa-cov provides ANSI SQL compliant API for accessing and analyzing of aligned sequencing reads data.

We present the first fully scalable, distributed, SQL-oriented solution designated for the depth of coverage calculations. SeQuiLa-cov, an extension to the recently released SeQuiLa [14] platform, runs a redesigned event-based algorithm for the distributed environment and provides convenient, SQL-compliant interface. The tool can be easily integrated with other applications implemented in Scala, R, or Python.

### Algorithm and implementation

#### Algorithm

Consider input data set, *read_set*, of aligned sequencing reads sorted by genomic position from a BAM file partitioned into the *n* data slices (*read_set_1_, read_set_2_,…, read_set_n_*) (Figure 1B).

In the most general case, the algorithm can be used in a distributed environment where each cluster node computes the coverage for the subset of data slices using the event-based method. Specifically, for the í-th partition containing the set of reads (*read_set_i_*), the set of *events_i,chr_* vectors (where *chr* is an index of genomic contig represented in *read_set*) is allocated and updated, based on the items from *read_set*. For all reads, the algorithm parses the CIGAR string and for each continuous alignment block characterized by *start* position and length *len* it increments by one the *events_i,chr_*(*start*) and decrements by one the value of *events_i,chr_*(*start + len*). To compute the partial coverage vector for partition *i* and contig *chr*, a vector value at the index *j* is calculated as follows:

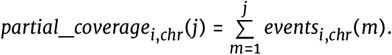

The result of this stage is a set of *partial_coverage_i,chr_* vectors distributed among the computation nodes. To calculate the final coverage for the whole *read_set*, an additional step of correction for overlaps between the partitions is required. An overlap *overlap_i,chr_* of length *l* between vectors *partial*_*coverage_i,chr_* and *partial_coverage_i+1,chr_* may occur on the partition boundaries where *l* tailing genomic positions of *partial*_*coverage_i,chr_* are the same as *l* heading genomic positions of *partial_coverage_i+1, chr_* (see Figure 1C).

If an overlap is identified then the coverage values from the *partial_coverage_i,chr_*’s *l*-length tail are added into the *partial_coverage_i+1,chr_*’s head and subsequently the last *l* elements of *partial_coverage_i,chr_* are removed. Once this correction step is completed, non-overlapping *coverage_i,chr_* vectors are collected and yield the final coverage values for the whole input *read_set*.

The main characteristic of the described algorithm is its ability to distribute data and calculations (such as BAM decompression and main coverage procedure) among the available computation nodes. Moreover, instead of simply performing full data reduction stage of the partial coverage vectors our solution minimizes required data shuffling among cluster nodes by limiting it to the overlapping part of coverage vectors. Importantly, SeQuiLa-cov computation model supports fine-grained parallelism at user-defined partition size in contrary to the traditional, coarse-grained parallelization strategies that involve splitting input data at a contig level.

#### Implementation

We have implemented SeQuiLa-cov in Scala programming language using the Apache Spark framework. To efficiently access the data from a BAM file we have prepared a custom data source using Data Source Application Programming Interface (API) exposed by SparkSQL. Performance of the read operation benefits from the Intel Genomics Kernel Library (GKL) [15] used for decompressing the BAM files chunks and from predicate pushdown mechanism that filters out data at the earliest stage.

The implementation of the core coverage calculation algorithm aimed at minimizing, whenever possible memory footprint by using parsimonious data types, e.g. *Short* type instead of *Integer*, and efficient memory allocation strategy for large data structures, e.g. favoring static *Arrays* over dynamic size *ArrayBuffers*. Additionally, to reduce the overhead of data shuffling between the worker nodes in the correction for the overlaps stage we used Spark’s shared variables [16] *accumulators* and *broadcast variables* (Figure 1C). Accumulator is used to gather information about the worker nodes’ coverage vector ranges and coverage vector tail values, that are subsequently read and processed by the driver. This information is then used to construct a broadcast variable distributed to the worker nodes in order to perform adequate trimming and summing operations on partial coverage vectors.

### Functionality

#### Supported coverage result types

SeQuiLa-cov features three distinct result types: *per-base, blocks*, and *fixed-length windows* coverage (Figure 1A). For *per-base*, the depth of coverage is calculated and returned for each genomic position making it the most verbose output option. The method producing block level coverage (*blocks*) involves merging adjacent genomic positions with equal coverage values into genomic intervals. As a consequence, fewer records than in case of *per-base* output type are generated with no information loss. The *fixed-length windows* the algorithm generates set of fixed length, tiling, non-overlapping genomic intervals and returns arithmetic mean of coverage values over positions within each window.

#### ANSI SQL compliance

SeQuiLa-cov solution promotes SQL as a data query and manipulation language in genomic analysis. Data flows are performed in SQL-like manner through the custom data source supporting convenient Create Table as Select and Insert as Select methods. SeQuiLa-cov provides a table abstraction over existing BAM/CRAM files, with no need of data conversion, which can be further conveniently queried and manipulated in a declarative way. The coverage calculation function *bdg_coverage*, as described in Algorithm sub-section, has been implemented as *table-valued* function(Figure 1D).

### Benchmarking

We have benchmarked SeQuiLa-cov solution with leading software for depth of coverage calculations, specifically samtools *depth*, bedtools *genomeCov*, sambamba *depth* and mosdepth (results of *DepthOfCOverage* from outdated GATK version are available in supplementary data). The tests were performed on the aligned WES and WGS reads from the NA12878 sample (see Methods for details) and aimed at calculating blocks and window coverage. To compare the performance and scalability of each solution, we have executed calculations for 1, 5, and 10 cores on a single computation node (see Table 2).

**Table 2.** Benchmarking leading solutions against SeQuiLa-cov on WES/WGS data in performing blocks and windows calculations

| data | operation type | cores | samtools | bedtools | sambamba | mosdepth | SeQuiLa-cov |
| --- | --- | --- | --- | --- | --- | --- | --- |
| WGS | blocks | 1 | 2h 14m 58s<sup>1</sup> | 10h 41m 27s | 2h 44m 0s | **1h 46m 27s** | 1h 47m 5s |
|  |  | 5 |  |  | 2h 47m 53s | 36m 13s | **26m 59s** |
|  |  | 10 |  |  | 2h 50m 47s | 34m 34s | **13m 54s** |
|  | fixed-length windows | 1 |  |  | 1h 46m 50s | **1h 22m 49s** | 1h 24m 8s |
|  |  | 5 |  |  | 1h 41m 23s | 20m 3s | **18m 43s** |
|  |  | 10 |  |  | 1h 50m 35s | 17m 49s | **9m 14s** |
| WES | blocks | 1 | 12m 26s<sup>1</sup> | 23m 25s | 25m 42s | **6m 43s** | 6m 54s |
|  |  | 5 |  |  | 25m 46s | 2m 25s | **1m 47s** |
|  |  | 10 |  |  | 25m 49s | 2m 20s | **1m 4s** |
|  | fixed-length windows | 1 |  |  | 14m 36s | **6m 11s** | 6m 29s |
|  |  | 5 |  |  | 14m 54s | 2m 8s | **1m 42s** |
|  |  | 10 |  |  | 14m 40s | 2m 14s | **1m 1s** |

Both samtools and bedtools calculate coverage using only a single thread, however, their results differ significantly, with samtools being around twice as fast. Sambamba positions itself as a multithreaded solution although our tests revealed that its execution time is nearly constant, regardless of the number of CPU cores used, and even twice as slow as samtools. Mosdepth achieved speedup against samtools in blocks coverage and against sambamba in windows coverage calculations, however, its scalability reaches limit at 5 CPU cores. Finally, SeQuiLa-cov, achieves nearly identical performance as mosdepth for the single core but the execution time decreases substantially for greater number of available computing resources which makes this solution the fastest when run on multiple cores and nodes.
<sup>1</sup>per-base results are treated as blocks output. Samtools lacks the functionality of blocks coverage calculations, however, we included this tool in our benchmark for completeness, treating its per-base results as blocks outcome assuming that both result types require nearly the same resources.

Samtools *depth* and bedtools *genomeCov* are both natively non-scalable and were run on a single thread only. Exome-wide calculations exceeded 10 minutes and genome-wide analyses took over two hours in case of samtools, while bedtools’ performance was significantly worse, i.e ~1.9x for WES and ~4.75x for WGS. Sambamba *depth* declares to take advantage of fully parallelized data processing with the use of multithreading. However, our results revealed that even when additional threads were used, the total execution time of coverage calculations remained nearly constant and greater than samtools’s result. Mosdepth shows significant speedup (~1.3x) against samtools when using single thread. This performance gain increases to ~3.7x when using 5 decompression threads, however, it does not benefit from adding additional CPU power. In case of fixed-length window coverage mosdepth achieves over ~1.3 speedup against sambamba.

SeQuiLa-cov achieves performance similar to mosdepth when run using a single core. However, SeQuiLa-cov is ~1.3x and ~2.5x as fast as mosdepth when using 5 and 10 CPU cores, respectively, demonstrates its better scalability. The similar performance characteristic is observed for both blocks and fixed-length windows methods.

To fully assess the scalability profile of our solution, we have performed additional tests in a cluster environment (see Methods for details). Our results show that when utilizing additional resources (i.e. more than 10 CPU cores), SeQuiLa-cov is able to reduce the total computation time to 15 seconds for WES and less than one minute for WGS data (Figure 2). Scalability limit is achieved for 200 and ~500 CPU cores in case of WES and WGS data, respectively.

**Figure 2.**
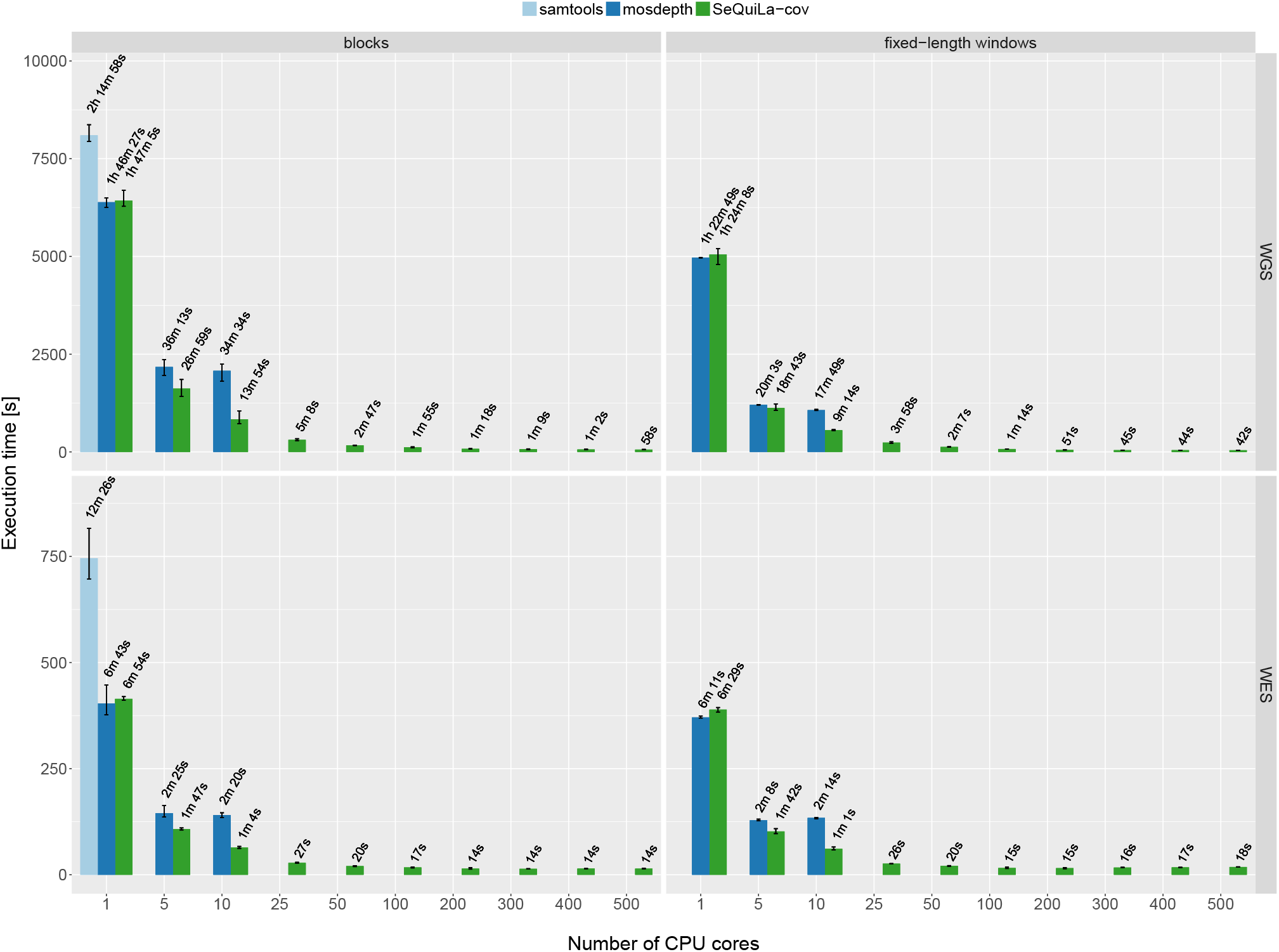
Performance and scalability comparison of samtools, mosdepth and SeQuiLa-cov. Each experiment setting was repeated several times and the height of each bar along with the corresponding error bars indicate the average, as well as the minimum and maximum execution time, respectively. The best pileup-based solution is definitely slower (two times for WGS calculations) than both event-based solutions what clearly shows the superiority of the latter one. Mosdepth execution time scales up to 5 cores, afterwards it shows no furthe gain in performance. SeQuiLa-cov has nearly the same execution time results as mosdepth for both blocks and windows calculations for a single core, but scales out desirably utilizing all 500 CPU cores on cluster nodes and at the same time performing WGS calculations in less than 1 minute.

To evaluate the impact of Intel GKL library on deflate operation (BAM bzgf block decompression), we have performed blocks coverage calculations on WES data on 50 CPU cores. The results showed on average ~1.18x speedup when running with Intel GKL deflate implementation.

Finally, our comprehensive functional unit testing showed that results calculated by SeQuiLa-cov and samtools *depth* are identical.

## Conclusions

The application of the recent advancements in big data technologies and distributed computing can contribute to both speeding up genomic data processing and management. Analysis of large genomic data sets require efficient, accurate, and scalable algorithms to perform calculations utilizing the computing power of multiple cluster nodes. In this work, we show that with sufficiently large cluster genome-wide coverage calculations may last less than a minute and at the same time being over 100x faster than the best single-threaded solution.

SeQuiLa-cov is one of the building blocks of SeQuiLa [14] ecosystem, which initiated the move towards efficient, distributed processing of genomic data and providing SQL-oriented API for convenient and elastic querying. We foresee that following this direction will enable the evolution of genomic data analysis from the file-oriented to table-oriented processing.

## Methods

### Test data

We have tested our solution using reads from NA12878 sample which were aligned to hg18 genome. WES data containing over 161 million of reads weights 17 GB and WGS data include over 2,6 billion of reads taking 272 GB of disk space. Both BAM files were compressed at the default BAM’s compression level (5).

### Testing environment

To perform comprehensive performance evaluation, we have setup a test cluster consisting of 28 Hadoop nodes (1 edge node, 3 master nodes and 24 data nodes) with Hortonworks Data Platform 3.0.1 installed. Each data node has 28 cores (56 with hyper-threading) and 512 GB of RAM, Yet Another Resource Negotiator (YARN) resource pool has been configured with 2640 virtual cores and 9671 GB RAM.

### Investigated solutions

In our benchmark we have used the most recent versions of the investigated tools i.e. samtools version 1.9, bedtools 2.27.0, sambamba 0.6.8, mosdepth version 0.2.3 and SeQuiLa-cov version 0.5.1.

### Availability of source code and requirements

- Project name: SeQuiLa-cov
- Project home page: http://biodatageeks.org/sequila/
- Source code repository: https://github.com/zsi-Bio/bdg-sequila
- Operating system: Platform independent
- Programming language: Scala
- Other requirements: Docker
- License: Apache License 2.0

### Availability of supporting data and materials

The Docker image is available at https://hub.docker.com/r/biodatageeks/. Supplementary information on benchmarking procedure as well as test data is publicly accessible at http://biodatageeks.org/sequila/benchmarking/benchmarking.html#depth-of-coverage.

## Declarations

### List of Abbreviations

API –: Application Programming Interface
BAM –: Binary Alignment Map
GKL –: Genomics Kernel Library
NGS –: Next Generation Sequencing
SQL –: Structured Query Language
YARN –: Yet Another Resource Negotiator
WES –: Whole Exome Sequencing
WGS –: Whole Genome Sequencing

## Consent for publication

Not applicable

### Funding

This work has been supported by the Polish budget funds for science in years 2016-2019 (Iuventus Plus grant IP2015 019874), as well as Polish National Science Center grant Preludium 2014/13/N/ST6/01843.

### Author’s Contributions

MW – conceptualization, formal analysis, investigation, software and writing. AS – data curation, formal analysis, investigation, software, visualization and writing. WK – formal analysis, investigation, writing. TG – formal analysis, supervision, investigation, visualization and writing. All authors approved the final manuscript.

